# *NvashA* function reveals temporal differences in neural subtype generation in cnidarians

**DOI:** 10.1101/2025.05.12.653478

**Authors:** Jamie A Havrilak, MingHe Cheng, Layla Al-Shaer, Whitney B Leach, Mia Yagodich, Dylan Faltine-Gonzalez, Michael J Layden

## Abstract

Understanding how cnidarians pattern their nervous systems can provide insight into the ancestral mechanisms of neurogenesis that are shared with bilaterians, shedding light on the evolution of nervous systems. While previous studies have revealed deeply conserved mechanisms for neural induction and progenitor selection between cnidarians and bilaterians, less is known about how distinct neuronal subtypes are specified over time in cnidarians. We utilized single-cell mRNA sequencing to profile *NvashA-*expressing cells across embryonic and planula-larva stages of *Nematostella* neurogenesis, and functional experiments identified a dynamic role for *NvashA* over time. Our analysis revealed that unique neuronal subtypes emerge at different developmental stages, providing evidence for temporal patterning in developing cnidarian nerve nets. This can provide a foundation to better our understanding of neurogenic gene regulatory networks, and to compare neurogenesis across cnidarians, and with bilaterians, to improve our knowledge of nervous system evolution.

## Introduction

Cnidarian and bilaterian nervous systems evolved from the nerve net found in their last common ancestor (Hejnol and Rentzsch, 2015; 2016). This shared ancestry has sparked an interest in studying cnidarian neurobiology to support comparative analyses with bilaterians and to provide insights into the evolution and function of nervous systems. Previous research has shown that both cnidarians and bilaterians use a conserved set of genes to regulate neurogenesis and pattern their nervous system (Chrysostomou et al., 2022; Faltine-Gonzalez et al., 2023; Kelava et al., 2015; Rentzsch et al., 2017). However, most cnidarian studies have focused on early embryonic stages or broadly acting neurogenic programs, and little is known about how specific neuronal subtypes are patterned during development. In this study, we aim to understand how neuronal subtypes emerge during embryonic and larval development, using the model cnidarian *Nematostella vectensis*.

In bilaterians, spatial and temporal cues work together to pattern neural subtypes (Doe, 2017; Erclik et al., 2017; Hartenstein and Stollewerk, 2015; Holguera and Desplan, 2018). Spatial cues confine specific neuronal fates to defined regions, while temporal cues within each region determine the birth order of distinct neuronal subtypes from a common progenitor. Temporal cues can also influence the identity of progenitors arising from a shared spatial domain. This strategy enables a limited pool of progenitors to sequentially produce neurons with distinct molecular identities, contributing to the complex architecture of bilaterian nervous systems; however, the evolutionary origins of temporal regulation of neurogenesis remain unresolved. Investigating whether similar mechanisms operate outside the bilaterian lineage is essential for understanding the ancestral strategies of nervous system patterning.

While the molecular details can differ between bilaterian species, general themes recur. Some subtypes arise from unique progenitor pools that share spatial locations but appear at different times (Batista-Brito et al., 2008; Sockanathan and Jessell, 1998). Others are produced in a fixed birth order, with each wave of progenitors inheriting the same spatial cues but unique temporal ones (Pearson and Doe, 2004). Or, the birth order of different subtypes that emerge from a single progenitor can be tightly regulated such that the order that different subtypes emerge is invariant (Kohwi and Doe, 2013; Okano and Temple, 2009; Pearson and Doe, 2004). These strategies - alone or in combination - generate neural diversity. For example, unipotent progenitors may emerge once from the neurogenic tissue at different times, each producing a specific subtype, thereby generating diversity (Pearson and Doe, 2004). Alternatively, multipotent progenitors could arise once and sequentially generate different subtypes (Doe, 1992; Telley et al., 2019). These mechanisms could operate together and maximize diversity from a limited pool of progenitors.

Neurogenesis in *Nematostella* begins during the blastula stage. Coordinated Mek and Notch signaling specify neuronal progenitors, which express the genes *NvsoxC, NvsoxB(2),* and *Nvath-like* (Layden et al., 2016; Layden and Martindale, 2014; Rentzsch et al., 2017; Richards and Rentzsch, 2014, 2015; Steger et al., 2022). As these progenitors differentiate into neurons, they express *NvashA*, although this expression does not persist in mature neurons (Layden et al., 2012; Rentzsch et al., 2017). Axial patterning programs impose spatial constraints on neurogenesis, restricting the emergence of different neuronal subtypes to specific regions along the oral-aboral axis during the gastrula stage (Faltine-Gonzalez et al., 2023). We hypothesize that neurogenesis continues beyond the gastrula stage, based on the presence of proliferating neuronal progenitors and the sustained expression of proneural *sox* and *ash* genes (Layden et al., 2012; Richards and Rentzsch, 2014; Steger et al., 2022). However, the progression of neurogenesis and the emergence of neuronal diversity beyond the gastrula stage remain unexplored.

Recent single-cell transcriptomic studies have revealed up to 35 neuronal cell states (Cole et al., 2024; Sebe-Pedros et al., 2018), although the exact number of fully distinct and differentiated neuronal types remains unknown. Further, we have not yet investigated in detail whether neuronal patterning in cnidarians includes a temporal component. In previous work, we tracked the emergence of neuronal subtypes labeled by the *NvLWamide::mcherry* reporter construct throughout larval stages (Havrilak et al., 2017), but this analysis did not clarify whether these subtypes are specified early and then mature at different rates, or if they arise at different times. Some cell types, such as cnidocytes, require days to mature. For instance, cnidocytes begin expressing subtype markers during gastrulation but do not reach morphological maturity until several days later in larval development (Babonis and Martindale, 2014; Marlow et al., 2012; Steger et al., 2022). Single-cell atlases have identified neuronal cell states specific to early or late developmental stages, yet they have not determined when the late-restricted cell states first appear (Steger et al., 2022) (Cole et al., 2024). The best-characterized neuronal subtype in *Nematostella,* the *Nvfoxq2d*+ neurons, arise throughout development from a unipotent progenitor specified at the gastrula stage (Busengdal and Rentzsch, 2017), but it is unclear if new progenitors emerge later in development.

Current studies do not clearly determine whether neural subtype specification in *Nematostella* occurs exclusively during gastrula stages with subsequent maturation during larval development, or whether new neuronal subtypes continue to emerge during larval stages. Because there is no existing method that allows for temporal lineage tracing of neuronal progenitors in *Nematostella*, we used an alternative strategy to infer temporal dynamics by examining *NvashA* expression and function across development. *NvashA* is expressed in newly born neurons during both early and late stages and regulates a set of target genes expressed throughout development. In some cases, these targets expand in cell number over time (Havrilak et al., 2021; Havrilak et al., 2017; Layden et al., 2016). Although *NvashA* clearly regulates targets across developmental stages, it remains uncertain whether it continues to regulate the same targets during later development. Notably, *NvashA* controls the differentiation of multiple neuronal subtypes (Layden et al., 2016), which suggests that we can use its expression to identify different neuronal subtypes arising at distinct time points.

Temporal patterning involves the specification of distinct neuronal fates during defined development windows. We hypothesized that if temporal patterning occurs in *Nematostella*, we would observe stage-specific shifts in *NvashA* targets or changes in the identity of newly born, maturing, and mature *NvashA^+^* cells. Our results show that *NvashA*-expressing cells are present in both early and late neuronal populations and that *NvashA* regulates unique target genes at the gastrula and mid-planula larva stages. These findings demonstrate that neural subtype specification is not limited to differential maturation. Instead, *Nematostella* continues to generate new neuronal fates throughout development. This work reveals dynamic temporal regulation of neural development in *Nematostella* and raises important questions about the evolutionary conservation of temporal patterning mechanisms.

## Results

### Single-cell atlas of *NvashA* expressing cells identifies embryonic and larval born neuronal subtypes

One indicator of temporal patterning during neurogenesis is a shift in the identities of neurons that are born at different times. To determine if distinct neurons are generated at different times during development in *Nematostella*, single-cell mRNA sequencing was conducted on gastrula through late-planula stage embryos (Supplemental Figure 1 and 2), and then *NvashA* positive cells from all time points were subset and clustered (Figure 1). *NvashA* was chosen because it is expressed following the cell’s terminal division (when immature neurons still express some neuronal progenitor markers) (Richards and Rentzsch, 2015) through maturation to distinct neuronal subtypes (Layden et al., 2012; Layden et al., 2016; Layden and Martindale, 2014). Thus, examining *NvashA*-expressing cells provides a window into neurons from a naive to fully differentiated state. In total, 23 cell state clusters representing neurons (fourteen clusters), cnidocytes (seven clusters), gland cells (one cluster), and early secretory cells (one cluster) were annotated (Figure 1). Clusters were named and assigned identities using previously described cell markers and nomenclature for neuroglandular and cnidocyte cell states when possible (Cole et al., 2024; Steger et al., 2022), or identities were assigned through differential gene expression analysis (Supplemental Figures 3-5, see Materials and Methods). For instance, the top differentially expressed genes (DEGs) for each cnidocyte state identified by Steger et al. (2022) were mapped onto our *NvashA* dataset to help determine cluster identities (Figure 1; Supplemental Figure 5).

**Figure 1:**
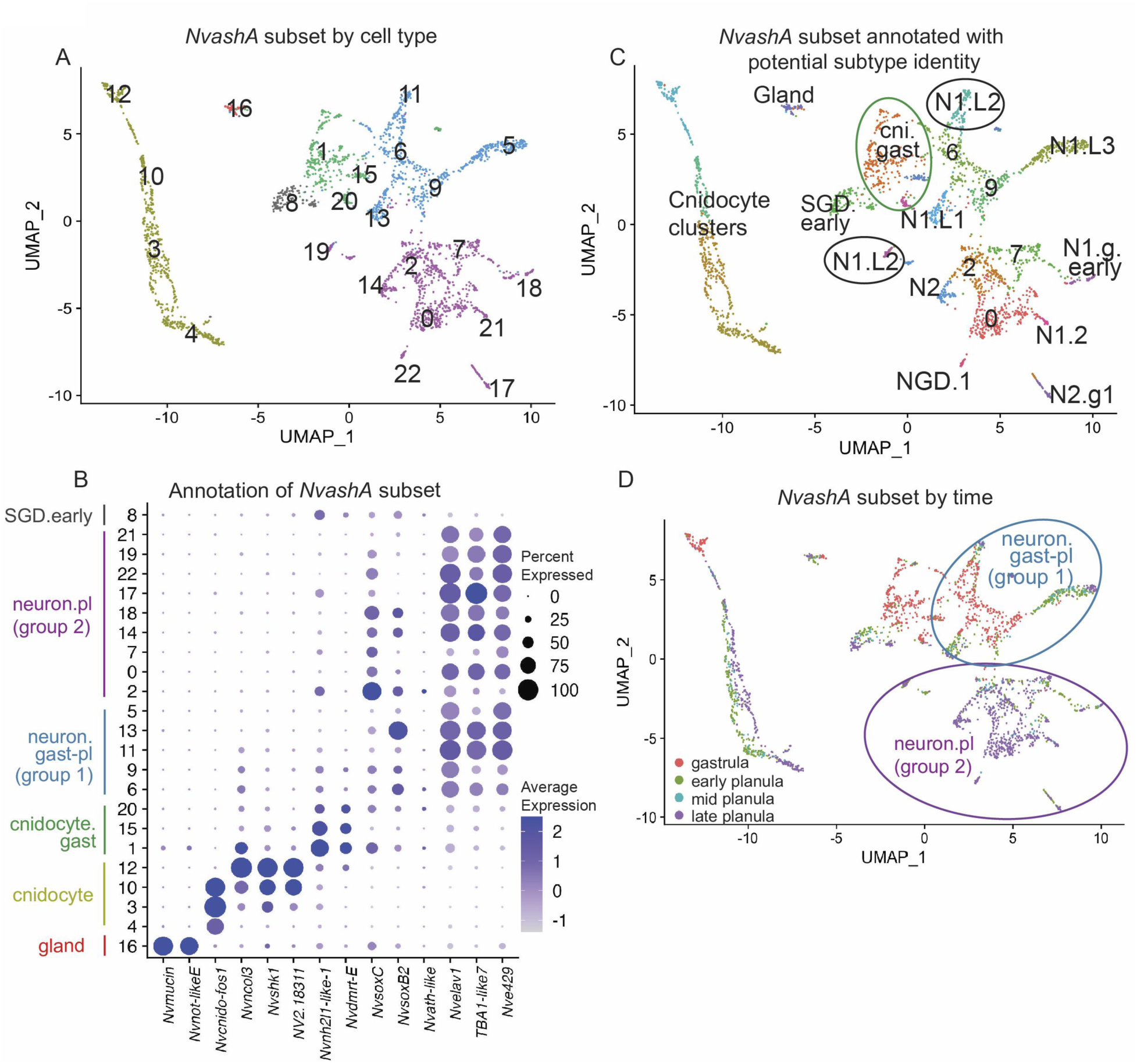
*NvashA* expressing cells comprise neural, cnidocyte, gland, and early secretory cell clusters. (A) UMAP of subclustered *NvashA* expressing cells from gastrula through late-planula stages. (B) Dotplot representation of the expression of the known marker genes used to annotate *NvashA* subset clusters in panel C. (C-D) UMAPs of *NvashA* subset, colored to identify the clusters by (C) subtype annotation and (D) developmental time point of origin. (D) Sub-clusters in group 1 (circled in blue) represent neural clusters containing cells from all four stages (neuron.gast-pl) while sub-clusters in group 2 (circled in purple) represent planula-stage only neural cells (neuron.pl).

**Figure 2:**
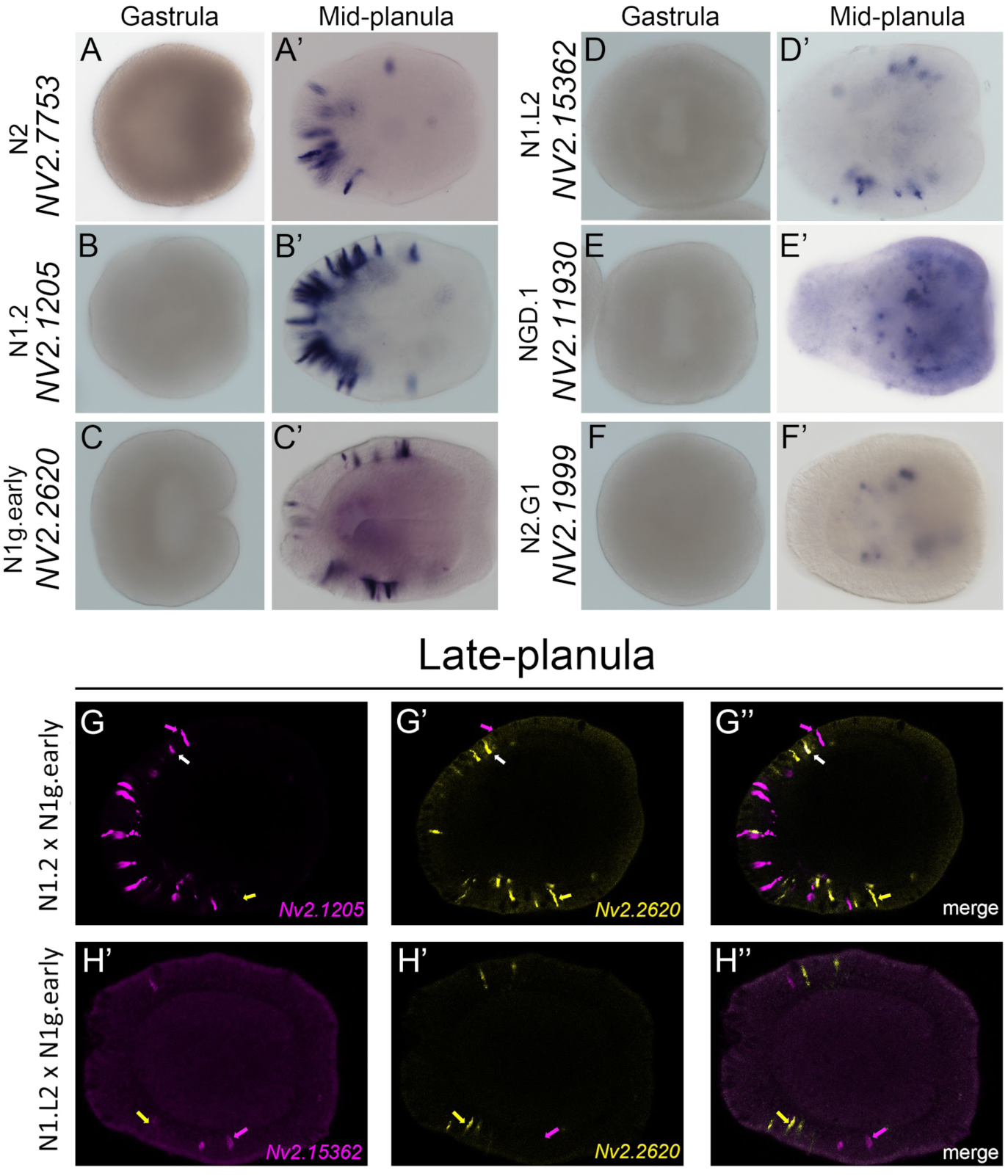
Larval *NvashA* expressing neural clusters likely represent distinct neuronal subtype markers. Expression of putative larval cluster marker genes at the gastrula (A-F) and mid-planula (A’-F’) stages. All markers showed no expression at the gastrula stage, which is consistent with the single-cell RNA sequencing data. At the mid-planula stage, different patterns of expression are shown via mRNA *in situ* hybridization. (A’-C’) *Nv2.7753*, *Nv2.1205*, and *Nv2.2620* showed an ectodermal salt and pepper pattern in the aboral and trunk domains. (D’-E’) *Nv2.15362* and *Nv2.11930* showed a salt and pepper expression pattern in the trunk domain, while (F’) Nv2.1999 is only expressed in the endoderm. (G-G’’) Double fluorescence mRNA *in situ* hybridization revealed that *Nv2.2620* partially overlaps with *Nv2.1205* in the trunk domain at the late-planula stage. Examples of *Nv2.1205* and *Nv2.2620* only expression are indicated with purple and yellow arrows respectively, while a single double positive cell is labeled with a white arrow. (H-H’’) No overlap was observed between *Nv2.2620* and *Nv2.15362* in late-planula animals. Oral end is positioned to the right.

Some cell states are present across all developmental stages surveyed, while others are more restricted (Figure 1C,D). Evidence of this is that some *NvashA*+ cnidocyte clusters were comprised of only early differentiating subtypes derived from gastrula stage embryos (‘cni.gast’), others were comprised of differentiating and mature cell state clusters from the three planula stages (‘cni.nem1’, ‘cni.maturing’, and ‘cni.mature’), and one cluster comprised cells from both gastrula and planula stages belonging solely to the nematocyte subtype lineage (‘cni.nem1’) (Figure 1A-C; Supplemental Figure 2G-J and Supplemental Figure 5). Further, these data are consistent with previous reports that cnidocytes can be detected by gene expression days before mature cnidocytes can be observed (Marlow et al., 2012). These findings suggest a potential role for *NvashA* during cnidocyte development, including the specification of the ‘cni.nem1’ nematocyte subtype (Steger et al., 2022) (Supplemental Figure 5).

### *NvashA* contributes to neuronal subtype diversity

To further characterize the neuronal cell states that are labeled by *NvashA,* DEGs that are uniquely over represented in each cell state were identified (Supplemental Figure 3A), and the top five DEGs were then mapped onto the neuroglandular atlas created by Cole et al. (2024) (Supplemental Figure 4A). Similarly, we performed the reciprocal analysis and mapped the top 5 DEGs from the Cole et al. (Cole et al., 2024) neural subtypes onto our *NvashA* dataset (Supplemental Figure 4B). This analysis determined that the N1.L1, N1.L2, N1.L3, N1.g.early, N1.2, NGD.1, and N2.g1 neural cell states are identifiable in our atlas (clusters 13, 11 + 19, 5, 18, 22, and 17, respectively) and also that a single *NvashA* cluster (cluster 14) mapped to several of the previously described N2 subtypes (N2.2-N2.4, henceforth grouped and called solely N2) (Figure 1D; Supplemental Figure 4). Cluster 8 is likely the SGD.early secretory cell type (Supplemental Figure 4). Five remaining clusters could not be given a specific cell state identity. Of those five, clusters 2, 6, 7, and 9 express progenitor markers (*NvsoxB(2), NvsoxC,* and/or Nv*ath-like*), and 2 and 7 also express less of the differentiated neuronal markers (*Nvelav1, NvTBA1-like7,* and/or *Nve429*) (Figure 1B and Supplemental Figure 2A-F). Because previous studies failed to detect proliferative *NvashA* positive cells (Richards and Rentzsch, 2014, 2015), we infer that clusters 2 and 7 are likely early differentiating post-mitotic neurons since they are not definitive progenitors, and do not have a signature of a specific neuronal type yet.

Interestingly, the neuronal clusters fall into two large groups; Group 1 contains six clusters that range from gastrula stage through late planula stages, and Group 2 contains eight clusters composed of cells from the larval stages only (Figure 1D; Supplemental Figure 2G-J).

Group 1 contains clusters 6 and 9 that appear to be immature neurons, as well as the previously identified neuronal subtypes N1.L1, N1.L2, and N1.L3 (Cole et al., 2024) (Figure 1C,D; Supplemental Figure 4). We also looked at the location of known neural markers and *NvashA* targets *NvLWamide-like, NvglraA-like, Nvserum amyloid A-like,Nvabcc4-like* (Supplemental Figure 6). We found that *NvLWamide-like* is expressed in N1.L1-N1.L3, while *Nvserum amyloid A-like* and Nvabcc4-like are only expressed in N1.L2 (Supplemental Figure 6). The N1.L1-N1.L3 clusters all contain cells from all four time points included in the atlas (Figure 1; Supplemental Figure 2). Interestingly the cells are arranged with the late-planula stage cells clustered away from the immature cells, potentially highlighting the fact that these cell types are born at gastrula stages, but take multiple days to fully mature. This is consistent with our previous characterization of *NvLWamide-like::mcherry* expressing cells that take multiple days to fully extend neurites (Havrilak et al., 2021).

Group 2 contains cluster 2, which has a gene expression profile consistent with early differentiating neurons (Figure 1B; Supplemental Figure 2). Stemming out from the immature cells we believe are present in clusters 7 and 0 are multiple clusters that have been previously identified as neuronal subtypes: N1.g.early, NGD.1, N1.2, N2.g1, and N2 (Cole et al., 2024) (Figure 1C; Supplemental Figure 4). To confirm that the larval neuronal clusters are late-born neurons, we cloned genes whose expression was enriched in those specific late neuronal clusters but not in any other cell type (see Materials and Methods; Supplemental Figures 3 and 4). Markers for the putatively later-born neurons were detectable by whole-mount mRNA *in situ* hybridization (ISH) at mid-planula but not gastrula stage embryos (Figure 2A-F’), and double fluorescent ISH confirmed that the expression of unique cell state markers are mostly non-overlapping (Figure 2G-H’’).

The subtypes that arise during gastrula stages (N1.L1-N1.L3) are clustered near cell states that appear to be newly post-mitotic (clusters 6 and 9), and some known neuronal subtype (N2 and N1.g.early) cluster near the planula-specific pool of newly postmitotic neuronal cell states (clusters 2 and 7). Further, markers of some of these late neural subtypes cannot be detected earlier in development (Figure 2). Thus, we argue that some neuronal fates are generated exclusively at the larval stage, indicating that a temporal component exists to pattern neural subtypes during *Nematostella* development. Clusters 6, 9 and 0 could potentially represent early differentiating versions (immature neurons) of the identified cell states along the trajectory of the identified cell states. For example, cluster 6 could be the early stages before the cells become like those represented in cluster 11, which we have identified as the N1.L2 subtype. However, a trajectory analysis needs to be performed to validate whether cluster 6 is a developmental precursor to N1.L2, for example.

### *NvashA* regulates different target genes at gastrula and planula stages

We previously demonstrated that *NvashA*, a proneural bHLH transcription factor, is necessary and sufficient for the specification of several neural genes, some of which have since been identified to represent distinct neuronal subtypes (Havrilak et al., 2021; Layden et al., 2012). Prior work, however, only examined the functional role of *NvashA* on the regulation of these genes at the early gastrula stage (∼24 hours post fertilization (hpf) (Layden et al., 2012; Layden et al. 2016). Because we know that neurogenesis occurs through the planula stages and continues into *Nematostella’s* adult life (Havrilak et al., 2021; Rentzsch et al., 2017) we sought to determine whether *NvashA* also regulates the specification of its known target genes at later stages. The impact of shRNA-mediated knockdown of *NvashA* on the expression of known targets was assessed by combining ISH and quantitative PCR approaches (qPCR) at embryonic gastrula and planula larva stages (Figure 3). Consistent with previous findings, the known early *NvashA* target genes *NvLWamide-like, Nvserum amyloid A-like, NvglrA3-like*, *and Nvabcc4-like* (Layden et al., 2012) had reduced expression following *NvashA* knockdown in gastrula stage embryos (Figure 3A-E). In mid-planula stage embryos, however, these same targets did not show reduced expression (Figure 3F-J’) despite the continued knockdown of *NvashA* (Figure 3F-F’). These results were validated by qPCR (Figure 3K), which suggests a potential differential temporal regulation by *NvashA* of these early targets, with *NvashA* no longer being required for their expression at later stages of development.

**Figure 3:**
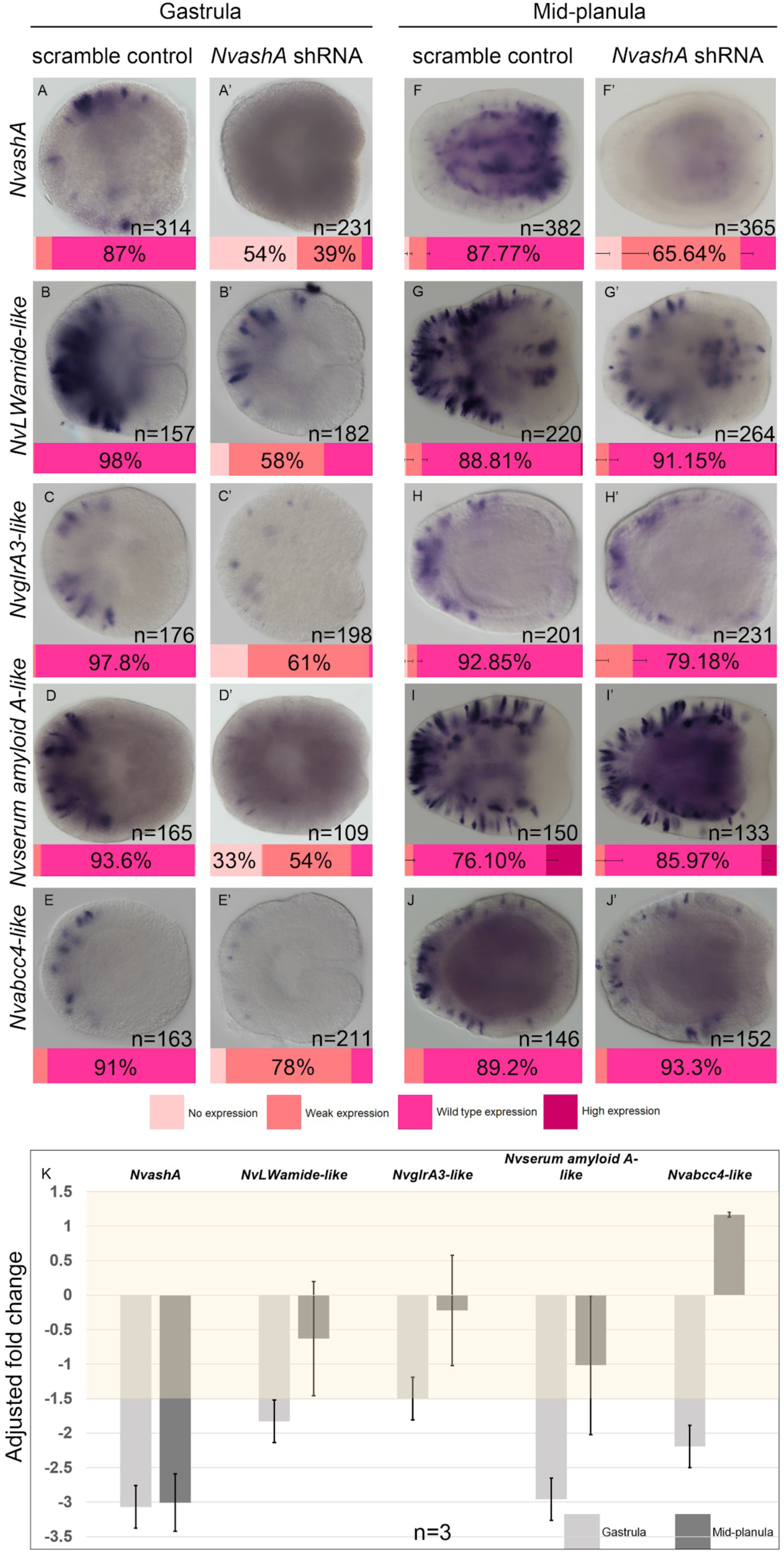
Temporal differences in *NvashA* regulation of known targets. (A-J’) mRNA *in situ* hybridization of *NvashA* and its known targets in control and *NvashA* shRNA injected embryos at (A-E’) gastrula and (F-J’) mid-planula stages. Images are lateral views with the oral pole oriented to the right. (K) Changes in gene expression of *NvashA* and its known targets were measured by qPCR at gastrula and planula stages by comparing the adjusted fold change between embryos injected with control and *NvashA* shRNA. An adjusted fold change of ± 1.5 was considered significant. The graph shows averages ± SE, *n* = 3 technical replicates.

Despite the sustained three-fold knockdown of *NvashA* observed by qPCR, we did see a slight recovery in the ISH signal of *NvashA* in a few cells in planula stage embryos, which suggests the knockdown of *NvashA* is weakened at later stages (Figure 3, compare A-A’ to F-F’). To determine if the lack of a phenotype at the mid-planula stage reflected reduced efficacy of the shRNA knockdown at later stages, we generated a *NvashA* null allele using CRISPR/Cas9 mediated gene editing. The functional bHLH domain of the *NvashA* coding sequence was replaced with a *Nvubiquitin::egfp* transgene (Supplemental Figure 7A). Only about 2% of offspring from wild-type crosses express no *Nvasha* throughout gastrula and planula stages (Supplemental Figure 7B). The offspring of heterozygous *NvashA^+/Ubi::eGFP-^*mutant crosses show no *NvashA* expression in approximately 27% of embryos from gastrula through planula stages, suggesting that homozygous *Nvasha^Ubi::eGFP-/Ubi::eGFP-^*mutant offspring are present at expected Mendelian ratios (Supplemental Figure 7B) and that the null mutation is not lethal at the planula stage (Supplemental Figure 7C-D, compare black lines 48-96 hpf). Therefore, we predicted that genes regulated solely by *NvashA* would not be expressed in roughly 25% of the mutant offspring population. To test this, we assessed the expression of the *NvashA* targets *Nvabcc4-like* and *Nvserum amyloid A-like* at gastrula and mid-planula stages. At the gastrula stage, expression of *Nvabcc4-like* and *Nvserum amyloid A-like* was reduced in 72.47% and 63.27%, respectively (Figure 4A’,B’), which is substantially more than what was classified as low expression in wild-type controls (*Nvabcc4-like*: 42.06%, *Nvserum amyloid A-like*: 24.82%) (Figure 4A,B). These results align with our expectation of 25% based on Mendelian ratios and are consistent with our observations following shRNA knockdown of *NvashA* at gastrula stages (Figure 3D-E’, K).

**Figure 4:**
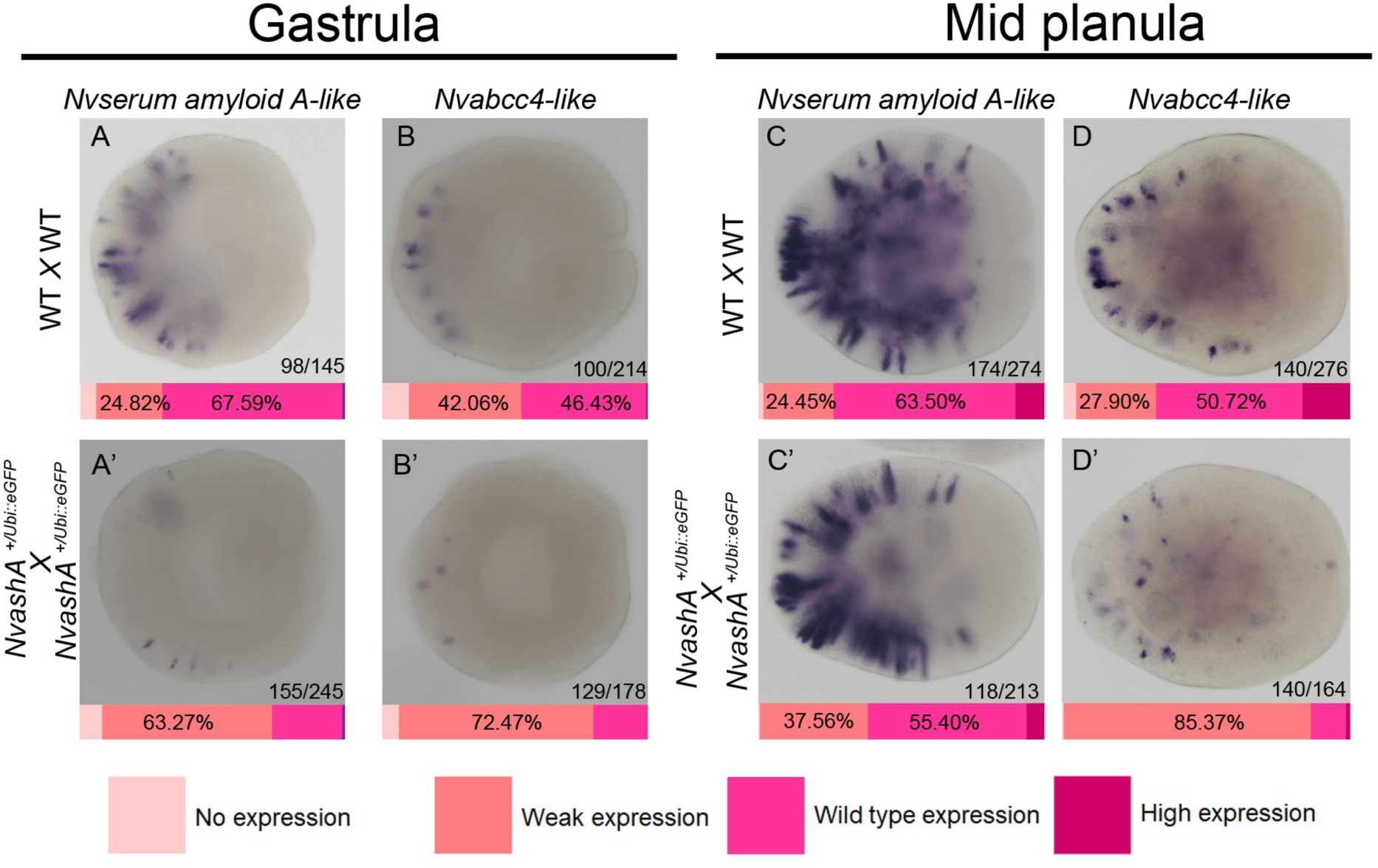
Nvabcc4-like and Nvserum amyloid A-like expression is unaffected in NvashA mutant offspring at mid-planula stage. Compared to wild-type, expression of *Nvserum amyloid A-like* is reduced in *NvashA^+/Ubi::eGFP^* X *NvashA^+/Ubi::eGFP^*mutant offspring at gastrula (A-A’) but not mid-planula stages (C-C’). Similarly, compared to wild-type offspring, *NvashA* mutant offspring exhibit reduced *Nvabcc4-like* expression at gastrula (B-B’) but not mid-planula stages (D-D’). Bars indicate the average percent of each phenotype observed in 1 technical replicate. The oral pole is oriented to the right in all images.

In contrast to our shRNA results, mutant offspring at the mid-planula stage display a reduction in *Nvabcc4-like* and *Nvserum amyloid A-like* expression (Figure 4C’,D’) when compared to wild-type controls of the same stage (Figure 4C,D). However, only a small proportion of the mutant offspring completely lacked *Nvabcc4-like* and *Nvserum amyloid A-like* expression (0.61% and 0.38%, respectively), but the number of embryos with weaker expression increased. Together, these data suggest that either *NvashA* expression is not required for all *Nvabcc4-like* or *Nvserum amyloid A-like* positive neurons, or that loss of *NvashA* results in a delay in their expression. The delay phenotype is difficult to test. Our strategy was to focus on the first 48 hours of development, when differentiated neuronal markers are being expressed for the first time. We fixed animals injected with *NvashA* shRNA and control animals injected with a scramble shRNA every six to ten hours beginning at 24 hpf and showed that expression of *Nvserum amyloid A-like* and *Nvabcc4-like* expression in *NvashA* shRNA knockdown animals increases over time by ISH, suggesting that in the absence of *NvashA,* differentiated neuronal markers are still expressed, but that their expression is delayed (Figure 5A-H’). This is despite mRNA expression of *Nvserum amyloid A-like* and *Nvabcc4-like* being significantly down in gastrula and mid-planula stage animals injected with *NvashA* shRNA compared to control animals by qPCR (Figure 5I-J).

**Figure 5:**
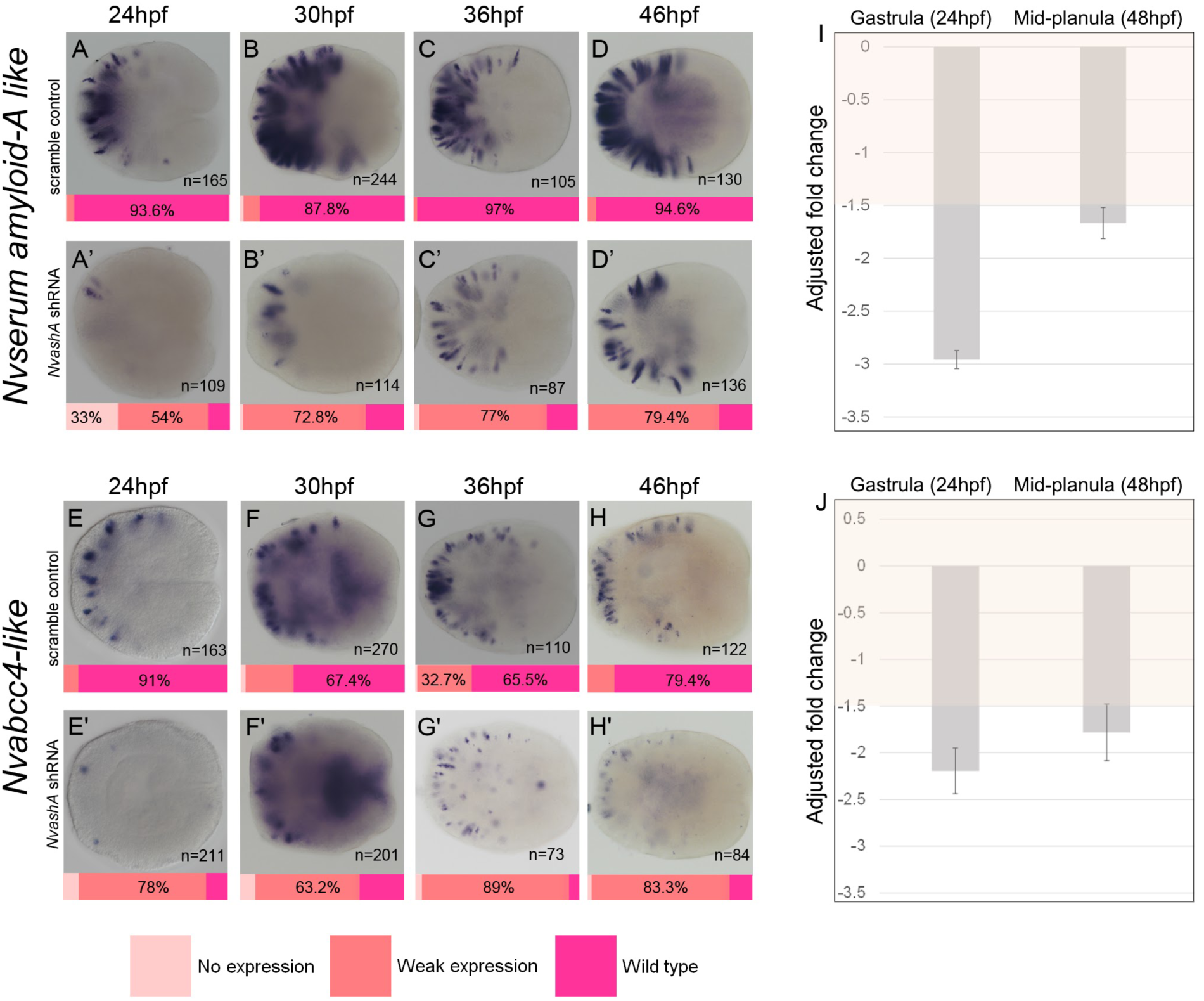
*NvashA* knockdown delays differentiation at the larval stage. mRNA *in situ* hybridization of *Nvserum amyloid-A like* (A-D’) and *Nvabcc4-like* (E-H’) at 24hpf, 30hpf, 36hpf, and 46hpf in embryos injected with *NvashA* shRNA or a scrambled control shRNA. At 24hpf, expression of both genes is reduced in *NvashA* knock down embryos (A-A’, F-F’). After 24hpf, expression of both *NvashA* knockdown embryos showed recovery and resembled the earlier stage of the control embryos (B-H’). qPCR results of *Nvserum amyloid-A like* (I) and *Nvabcc4-like* (J) at 24hpf, and 48hpf. (A-H’) Bars indicate the average percent of each phenotype, and in (I-J) bars show averages ± SE, *n* = 3. Oral pole is oriented to the right.

To further determine if *NvashA* is necessary for the specification of late-born neural subtypes, we assessed the expression of the late putative neural subtype marker genes *Nv2.2620* and *Nv2.1206* in planula-stage *NvashA* mutant embryos (Figure 6). In both cases, we observed an increase in the number of embryos that were weakly expressing the genes when heterozygous *NvashA^+/Ubiq::eGFP-^* mutants were crossed when compared to wild-type crosses. This suggests that the mutants also show a reduction of late *NvashA* cell state markers. However, we never observed a total loss of expression in 25% of the embryos, which is what would be expected if loss followed Mendelian ratios. Additionally, we saw substantial variability in the decrease in expression of *Nv2.2620* between two replicate experiments (Figure 6A-A’’, note error bars). Because *Nematostella* do not develop synchronously, and subtle temperature variations can occur with increased access to the incubator, we suspect that the variable phenotypes also reflect a delayed phenotype in later expressed genes. Regardless, unique later expressed neuronal markers exist and their expression is influenced by *NvashA*.

**Figure 6:**
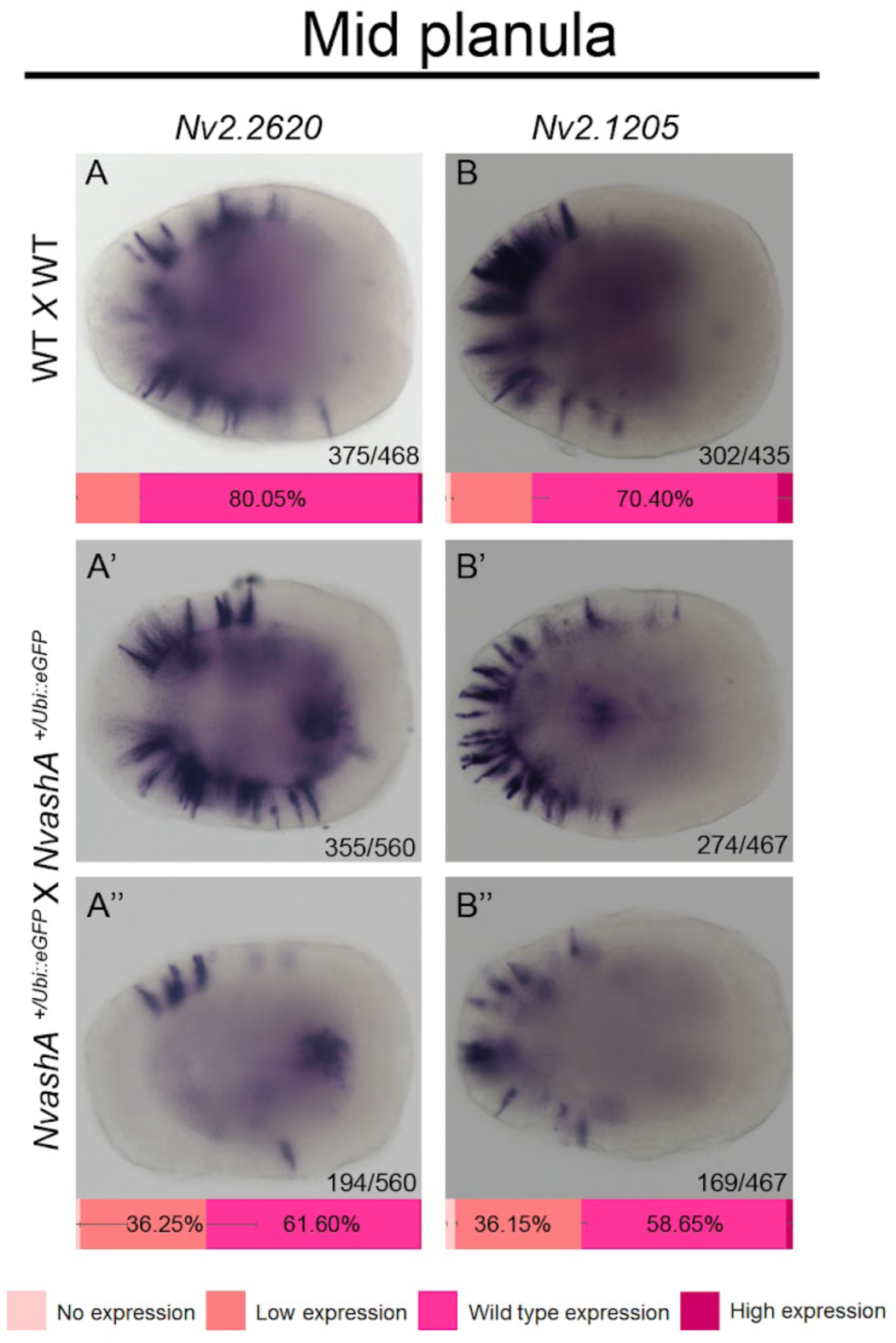
*NvashA* contributes to the specification of late-born neural subtypes. Expression of *Nv2.2620* and *Nv2.1206* was compared between mid-planula stage offspring resulting from (A-B) wild type or (A’-B’’) *NvashA* heterozygous mutant crosses using mRNA *in situ* hybridization. For both (A’-A’’) *Nv2.2620* and (B’-B’’) *Nv2.1206,* there was an increased proportion of heterozygous mutant offspring with weak expression compared to control offspring. Bars indicate the average percent of each phenotype ± SE, *n* = 2 technical replicates. Oral pole is oriented to the right.

The identification of distinct populations of neurons at embryonic and larval stages, and the dynamic role of *NvashA* throughout, suggests that neuronal diversity in the *Nematostella* nerve net relies on spatially localized signals (Faltine-Gonzalez et al., 2023; Nakanishi et al., 2012; Rentzsch et al., 2017) as well as temporally regulated mechanisms, and provides a foundation for dissecting gene regulatory logic of temporal fate specification in early-diverging animals.

## Discussion

### Temporal patterning contributes to neuronal diversification in *Nematostella*

Our results demonstrate that *NvashA* regulates the generation of distinct neuronal subtypes at different developmental stages, reflecting a dynamic neurogenic program in *Nematostella*. This provides the first clear evidence of temporal specification of neural subtypes in cnidarians and suggests that mechanisms for temporally controlled neural diversification may have emerged before the evolution of bilaterians, offering a new perspective on how nervous system complexity originated.

Although prior studies have proposed a temporal component to neurogenesis in *Nematostella*, they have not characterized temporal patterning of neural subtypes. For example, neurogenesis begins in the aboral ectoderm during early development and later occurs in the oral region and the endoderm (Nakanishi et al., 2012; Rentzsch et al., 2017). Additionally, spatial cues along the oral-aboral axis influence neuronal subtype identity at early stages (Faltine-Gonzalez et al., 2023). However, these studies focused on early developmental windows and did not investigate the timing of subtype specification. Here, we show that temporal cues contribute to the generation of distinct neural subtypes in *Nematostella*. Using a single-cell atlas of *NvashA*-expressing cells from gastrula through late planula stages, we identified two temporally distinct neural populations: One group emerges during the gastrula stage and differentiates into established subtypes, while a second group arises during planula stages and differentiates into different previously undescribed subtypes.

The specification of neuronal subtypes follows a tightly orchestrated process, with most bilaterian nervous systems patterned through spatial and temporal cues (Doe, 2017; Erclik et al., 2017; Hartenstein and Stollewerk, 2015; Holguera and Desplan, 2018). In *Nematostella*, spatial patterning along the oral-aboral axis defines where specific neural subtypes form during gastrula stages (Faltine-Gonzalez et al., 2023). However, the contribution of temporal cues has remained largely unexplored. Our findings support a model in which the timing of progenitor birth also influences neuronal identity. *NvsoxB(2)*^+^ progenitors display limited proliferation, and currently *Nvfoxq2d*^+^ cells are the only known unipotent progenitors (Busengdal and Rentzsch, 2017). Although *NvsoxC*^+^ cells have been proposed as multipotent proliferative stem cells (Steger et al., 2022) more data are needed to confirm this. Our data suggests that temporal regulation, rather than extensive proliferation of multiple unipotent progenitors, drives subtype diversification in *Nematostella*.

### NvashA’s dynamic role across development

*Nvasha* is a well-studied neurogenic transcription factor critical for early neurogenesis (Layden et al., 2012; Layden et al., 2016; Layden and Martindale, 2014; Richards and Rentzsch, 2015). Consistent with this, 14 cell states in our *NvashA*-specific atlas were classified as neural. However, we observed that knocking down *NvashA* had minimal impact on neural marker gene expression at planula stages. This suggests that *NvashA’s* function shifts over time. It likely activates neural transcription during early development and later contributes to neural maturation, possibly in combination with stage-specific regulators. Although, this remains to be investigated. Additional evidence of its dynamic function comes from the failure of *NvashA^Ubiq::eGFP-/Ubiq::eGFP-^* mutants to reach the adult polyp stage (Supplemental Figure 7C,D), indicating broader physiological defects linked to disrupted nervous system development. These findings underscore the importance of both early and late functions of *NvashA,* which likely operate through distinct gene regulatory networks.

### Implications for nervous system evolution

Our data suggest that temporal patterning is an ancient and conserved feature of neurogenesis. The presence of time-restricted waves of neurogenesis in a cnidarian implies that temporal mechanisms evolved prior to the central nervous system. Studying both spatial and temporal patterning strategies across early-diverging metazoa may offer further insights into the origins of neural diversity.

The next critical step involves identifying the molecular cues that mediate temporal patterning in *Nematostella* and linking gene regulatory networks back to specific neural subtypes. Future questions should include which transcription factors or signaling pathways show temporal restriction, how do they integrate with spatial cues to specify neuronal subtype identity, and whether the spatial cues that pattern neurons during gastrula stages continue to influence neuronal identity at later stages? Answering these questions will require time-resolved lineage tracing of neural progenitors, and inducible tools for investigating the functional roles of potential important factors at larval through polyp stages.

## Conclusion

Our single-cell transcriptomic analysis of *NvashA* expressing cells during embryonic and larval development reveals that temporal patterning plays a key role in specifying neuronal subtype identity in *Nematostella*. Many subtype-specific neural markers initiate *de-novo* at larval stages, and distinct neuronal subtypes arise in a time-dependent manner. Functional perturbation of *NvashA* further reveals that its role is both subtype- and developmental stage- specific. These results suggest that cnidaria use both spatial and temporal mechanisms to drive neural diversification. Our findings demonstrate that temporal patterning is an evolutionarily conserved strategy for generating neuronal diversity. Future work will aim to identify the upstream temporal cues and downstream effector molecules that together define specific neural trajectories throughout *Nematostella* neurogenesis.

## Materials and Methods

### Animal Care and microinjection

Adult *Nematostella* were maintained in 11-13 parts per thousand *Nematostella* Medium (NM) (Tropic Marin Pro-Reef Sea Salt), pH 8.1-8.2, in the dark at 17°C, given weekly water changes, and fed *Artemia sp.* four times per week. Spawning was induced as previously described (Havrilak and Layden, 2019). Microinjections were performed on wild type embryos at the single-cell stage as previously described (Havrilak and Layden, 2019; Layden et al., 2013). Embryos were raised at 24°C to the desired stage in the dark (Layden et al., 2013). Before fixation embryos were cleaned and checked for normal morphology, and planula were relaxed with 7.14% MgCl_2_.

### Colorimetric *in situ* hybridization

Embryos were fixed and stored as previously described (Havrilak and Layden, 2019; Wolenski et al., 2013). Colorimetric *in situ* hybridization experiments were performed using previously published methods (Wolenski et al., 2013). Probe primers used in this study can be found in Supplemental Table 1. Quantifications were performed by counting the number of embryos in the sample with wild type, high, weak, or no expression for each gene as viewed under a dissection microscope (Nikon SMZ1270). Afterwards, embryos were placed in glycerol and mounted for DIC imaging using a Nikon NTi compound microscope with a Nikon DS-Ri2 color camera and Nikon Elements software. Gene-specific digoxigenin riboprobes were prepared as previously described (Wolenski et al., 2013). Probes were denatured (80°C for 10 min), then snap cooled (ice 2 min), then brought to hybridization temperature (60°C for 10 min) before they were added to the samples.

### Double Fluorescent *in-situ* hybridization

Samples were photobleached with 3% H_2_O_2_ in MeOH under a light box (1 hr), rehydrated through 75%, 50%, and 25% MeOH in PBSTw and washed in PBSTw (5x5 min). They were treated with 60 µg/µL proteinase K (2 min), post-fixed in 4% paraformaldehyde in PBSTw (1hr), and washed again in PBSTw (5x5 min). Samples were then incubated in 50:50 PBST with pre-hybridization buffer (10 min, room temperature (RT)), then full pre-hybridization buffer (10 min, RT), followed by overnight incubation in hybridization buffer at 60°C overnight. The next day, we diluted gene-specific digoxigenin and fluorescence riboprobes (Supplemental Table 1) 1:1 in hybridization buffer (1.0 ng/µL), denatured them at 80°C for 10 minutes, snap-cooled on ice for two minutes, then incubated them at 60°C for 10 minutes. We hybridized samples at 60°C for 60 hours. After probe recovery, we washed samples in full hybridization buffer (10 min), then twice in pre-hybridization buffer (20 min ea., 60°C). We stepped samples into 2X SCC through 25%, 50%, and 75% pre-hybridization buffer (30 min ea.), then incubated them in 100% 2X SSC (30 min ea., 60°C), followed by three 0.1X SSC washes (20 min ea., 60°C). Next, we stepped samples into TNT through 75%, 50%, and 25% 0.1X SSC in TNT (5 min ea., RT). We blocked samples in TNT + 5% sheep serum (Sigma, S3772) + 1% Roche blocking reagent (11096176001) for 2 hours while rocking at RT, then incubated them overnight at 4°C with anti-digoxigenin-POD Fab fragments (Roche, 11207733910) in blocking buffer at [1:500] on a shaker. The next day, we washed samples eight times in TNT (20 min ea., RT, rocking), developed the digoxigenin signal using Cy3-TSA (AbexBio #K1051), and quenched POD activity with three TNT/0.3% H_2_O_2_ washes in the dark without rocking. We blocked again in TNT + 5% sheep serum + 1% Roche blocker (2 hrs, RT, rocking), then incubated overnight at 4°C with anti-fluorescein-POD Fab fragments (Roche, 11426346910) at [1:500] in blocking buffer. We washed samples eight times in TNT (20 min ea., RT, rocking), developed the fluorescein signal with HyperFluor 488 (AbexBio #4301) TNT washes (10 min ea.), followed by PBSTw washes (6x20 min, RT). Samples were washed in fresh PBSTw overnight (covered), incubated in Scale A2 at 4°C, then mounted and imaged on a Zeiss LSM 800 confocal microscope. Images were processed using Imaris (Bitplane) imaging software.

### Generation and genotyping of *NvashA* null mutant

The *NvashA* mutant was generated using CRISPR/Cas9 following a previously published protocol for *Nematostella (Ikmi et al., 2014)*, except that homologous recombination was directed to occur within the *NvashA* locus. Two guide RNAs (gRNAs) were generated to target PAM sites in the *NvashA* locus (Supplemental Table 4), one in the 5’ UTR (-33bp) and the other at the end of exon 1 after the conserved bHLH domain, (+395bp) (Supplemental Figure 7A). gRNAs were designed, cloned, and synthesized as previously described (Ikmi et al., 2014). The donor vector includes a *Nvubiquitin* enhancer upstream of the enhanced green fluorescent protein (eGFP) coding sequence, which together are flanked by 1Kb long homology arms (HA) that match the *NvashA* locus sequence up and downstream from the two PAM sites (Supplemental Figure 7A, Supplemental Table 1). The donor vector was generated by modifying the vector from Ikmi et al. (2014) to replace the existing enhancer and HA sequences. Microinjection was used to introduce a mixture of both gRNAs, Cas9 enzyme, and donor vector plasmid into wild-type embryos. Mutants were confirmed by PCR genotyping using two primer sets that share the same forward primer (upstream of *NvashA* exon 1) but have different reverse primers that match the sequence in *NvashA* exon 1 or the *Nvubiquitin* enhancer (Supplemental Figure 7E, Supplemental Table 1). In combination, the presence or absence of bands using these two primer sets can be used to distinguish between individuals with the following genotypes: *NvashA^+/+^, NvashA^+/Ubiq::eGFP-^,* and *NvashA^Ubiq::eGFP-/Ubiq::eGFP-^*(Supplemental Figure 7E).

DNA for genotyping was collected from either whole animal or tissue fragment that was treated with 200µl of 100% EtOH for 5min, then the EtOH was removed, and the sample was air dried, then treated with 50µl of 50mM NaOH for 30min at 95°C. This was followed by neutralization by adding 5µL of 1M Tris, pH8.0.

### shRNA preparation and delivery

shRNAs were designed and synthesized as described (Ikmi et al., 2014; Karabulut et al., 2019), and stored at −80°C in single-use aliquots. To achieve a sufficient knockdown, a combination of 2 shRNAs targeting *NvashA* were combined (500 ng/µl of each) for a final concentration of 1000 ng/µl. A previously published scrambled sequence shRNA was used as the control for all shRNA injections (Karabulut et al., 2019) and prepared to a total concentration of 1000 ng/µl. Similar volumes of shRNA were injected for control and *NvashA*-knockdown groups. The primers used to synthesize the two shRNA sequences targeting *NvashA* in this study can be found in Supplemental Table 1.

### Quantitative PCR

For qPCR experiments, total RNA was extracted from embryos at several developmental stages using phenol-chloroform phase separation and isopropanol precipitation as previously described (Röttinger et al., 2012). Assessment of gene knockdown via qPCR, including cDNA generation, was performed using a poly dT primer for first-strand synthesis as previously described (Layden et al., 2016). qPCR primers used in this study were previously published (Layden et al., 2012).

### Cell Dissociation and FACS

Dissociation protocol was modified from Torres-Mendez et al. (2019), and all solutions were filter sterilized with 0.2 µm syringe filters (VWR) before use. *Nematostella* embryos at the desired developmental stages were collected in 15 mL falcon tubes. After removal of NM, embryos were acclimated into calcium and magnesium free *Nematostella* media (CMF/NM, pH ∼8) for 5 minutes at RT, then washed in CMF/NM + EDTA for 10 minutes at RT. After removal of all but 0.5 ml of CMF/NM + EDTA, 4.5 mL of pre-warmed CMF/NM + EDTA + 0.25% Trypsin (pH 7.4-7.6) was added, for a total of 5 mL volume, and then samples were placed at 37°C for enzymatic digestion. The tissue was gently pipetted up and down ∼10x every 5-10 minutes with a p1000 pipette tip to help dissociate to single cells. Total time for enzymatic and mechanical dissociation was between 20-40 minutes depending on the stage of the animals. After cells were fully dissociated, 5 mL (or equal vol.) of cold CMF/NM + 0.5% BSA (pH 7.8) was added to stop the digestion and samples were kept on ice. Cells were then filtered through a 30 µm filter (CellTrics) and spun at 800 g for 10 minutes at 4°C to pellet the cells. Cells were washed with cold CMF/NM + 0.5% BSA and pipetted gently up and down with a cut pipette tip, spun again at 800 g for 7 minutes at 4°C, and resuspended in 2 mL cold CMF/NM + 0.5% BSA.

To assess cell viability and to allow for acquisition of live cells and exclusion of dead cells during the sort, cells were stained with Hoechst 33342 (2 drops/mL, incubated 15-30 min on ice in the dark) and Sytox green (1 µL/mL, incubated 20 min on ice in the dark). A subset of the cells were aliquoted into 1.5 mL tubes and stained with either Hoechst 33342 as a live cell-labeled control, or microwaved for 10 seconds (in an open tube) and stained with Sytox green as a dead cell-labeled control, or left unstained as a negative control for both stains.

Sorting of cells from early- mid-, and late-planula stage animals was performed on a Sony SH800S. The machine and collection chamber were pre-chilled to 4°C prior to sorting. We sorted and collected 100,000 Hoechst positive/Sytox negative cells into a 1.5mL Eppendorf tube (excluding also potential doublets and debris). After sorting, cells were spun at 600 g for 5 minutes at 4°C and resuspended to a targeted single cell suspension between 700-1200 cells/µl (1000 cells/µl optimal) in CMF/NM + 0.5% BSA. Cell concentration and viability were determined by visual inspection on a hemocytometer. Four, two, one, and two libraries were generated and sequenced for gastrula (24 hpf), early- (48 hpf), mid- (72 hpf), and late-planula (96 hpf) stages grown at 22°C, respectively.

Sequencing data from the same developmental stages were aggregated using the merge function during Seurat analysis. Data from gastrula-stage animals were previously published (Faltine-Gonzalez et al., 2023) and reanalyzed for this study.

### Single-cell RNA sequencing and analysis

The 10X genomics protocol was performed according to the manufacturer’s instructions for the Chromium Next GEM single cell 3’ Reagents Kits v3.1 (Dual Index). To prepare the single cell suspension for GEM generation and barcoding, a single cell master mix was assembled for a targeted cell recovery of 10,000 cells and loaded onto a 10X Genomics Chromium Chip, then run on a 10X Chromium Controller. Post GEM-RT cleanup, cDNA amplification, and library construction were conducted according to the manufacturer’s protocol. Library quality was assessed on an Agilent bioanalyzer. Illumina sequencing and processing of the raw data through Cell Ranger were performed by the Johns Hopkins Single Cell and Transcriptomics Core. Raw sequencing processed in Cell Ranger was aligned to the *Nematostella vectensis* genome (Zimmermann et al., 2023), and filtered count matrices were used for downstream analysis.

All downstream analyses were performed in R/RStudio. Low-quality cells were identified as cells containing > 10% mitochondrial counts, or cells expressing < 250 or >10000 genes, or cells containing < 500 or > 50000 unique molecular identifiers, and those cells that failed to meet quality control parameters were excluded. Potential doublets were removed with DoubletFinder (McGinnis et al., 2019). Filtered datasets were analyzed using standard Seurat pipelines (Hao et al., 2021) and modifications of previous publications (Faltine-Gonzalez et al., 2023; Steger et al., 2022). Clusters were determined and visualized with runUMAP (Uniform Manifold Approximation and Projection) and DimPlot functions in Seurat. Clusters were then annotated by plotting the expression levels of known cell type marker genes using Seurat DotPlot and FeaturePlot functions (Cole et al., 2024; Steger et al., 2022) (Supplemental Figures 3-8; Figure 4B). Markers used in this study for the annotation of clusters: Gland cell markers: *Nvmucin* and *Nvnot- likeE*; Cnidocytes: *Nvcnido-fos1*, *Nvncol3;* Neural Progenitor Cells: *NvsoxC*, *NvsoxB(2)*, *Nvath-like*; Neurons: *Nvelav1*, *NvTBA-like1*, *Nve490*; Pharyngeal Ectoderm; *NvfoxA*; Mesendoderm; *NvsnailA*; Trunk/Aboral Ectoderm; *Nvwnt2*, *NvSix3/6* (Cole et al., 2024; Steger et al., 2022) (see Supplemental Table 1 for gene IDs).

To investigate only cells that express *NvashA*, the subset function in Seurat was used on each developmental stage dataset to subset cells based on their expression level of *NvashA*. The *NvashA* subset datasets from each developmental time point were then combined using the merge function and reanalyzed with the standard Seurat pipelines.

Cluster-specific markers from the combined *NvashA* subset were computed using the FindAllMarkers function. Mapping the expression of the top 5 markers from each neural cluster back onto the *NvashA* dataset allowed for the identification of those clusters with differentially expressed genes, which we hypothesized could represent markers for those potential unique cell states (Supplemental Figure 3A). Clusters 11, 13, 5, 7, 14, 18, 17, 22, 19, and 21 had specific expression of their top markers to their own cluster (Supplemental Figure 3A) and restricted expression in neural cells (Supplemental Figure 3B), identifying them as potentially representing neuronal subtypes. Clusters with markers that were not restricted to their respective cluster (0, 2, 6, and 9) were excluded from later neuronal subtype analyses. Sub-annotations of cnidocytes and neurons into particular subtypes with the *NvashA* subset was accomplished in two ways: 1) by mapping the identified cluster markers onto the Cole et al. (2024) neuroglandular single-cell atlas/dataset (Supplemental Figure 4A), and 2) by assessing whether our clusters were enriched for genes found to be to enriched in the previously identified neural and cnidocyte subtypes (Cole et al., 2024; Steger et al., 2022) (Supplemental Figures 4B and 5A).

## CrediT authorship contribution statement

**Jamie Havrilak:** Conceptualization, Methodology, Software, Validation, Formal analysis, Investigation, Data curation, Writing- original draft, Writing- review and editing, Visualization

**MingHe Cheng:** Conceptualization, Methodology, Formal analysis, Investigation, Writing- original draft, Visualization

**Layla Al–Shaer:** Methodology, Software, Validation, Formal analysis, Investigation, Data curation, Writing- review and editing, Visualization

**Whitney Leach:** Methodology, Validation, Formal analysis, Investigation, Writing- review and editing

**Mia Yagodich:** Formal analysis, Investigation

**Dylan Faltine-Gonzalez:** Conceptualization, Methodology, Investigation

**Michael J Layden:** Conceptualization, Methodology, Writing- original draft, Writing- review and editing, Supervision, Project administration, Funding acquisition

## Acknowledgments

We thank David Balli for his help with bioinformatic and computational support, and Nolan Jetter for his technical assistance. This work was funded by the Department of Health and Human Services National Institute of General Medical Sciences (R01GM127615) and the National Science Foundation (NSF CAREER 1942777).

---

**Supplemental Figure 1:**
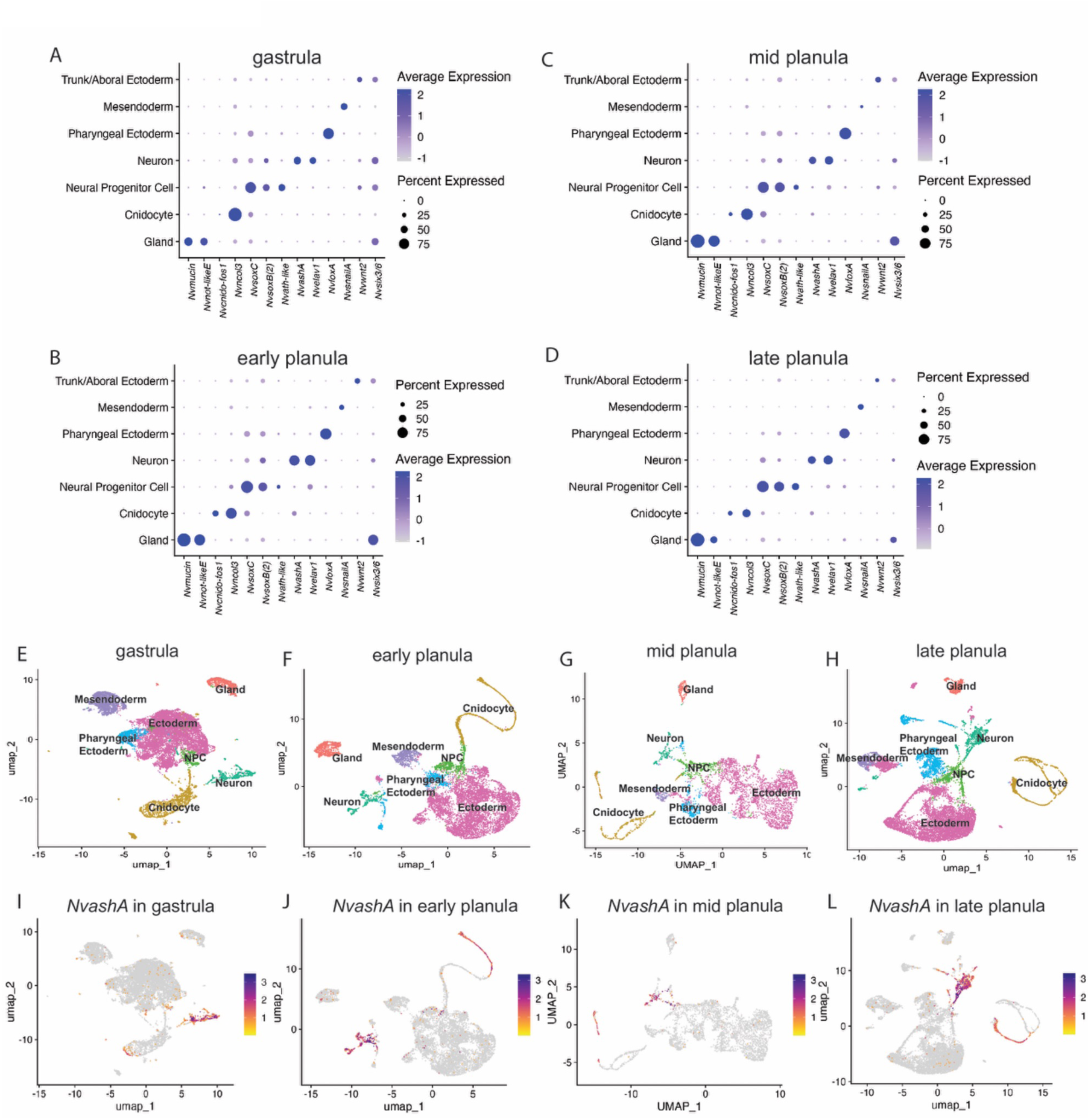
Single-cell RNA sequencing of gastrula through late-planula stages reveals *NvashA*-positive neural, cnidocyte, and glandular cell clusters. (A-D) Dot plot illustrations of the expression of marker genes utilized for annotating the clusters constructed for each developmental stage from which scRNA sequencing data was obtained. The size and intensity of a dot represent the number of cells that express that gene and the relative expression levels respectively. (E-H) UMAP of each time-point examined in this study, illustrating the annotated clustering of cells. (I-L) UMAPs highlighting cells that express *NvashA* within each time-point.

**Supplemental Figure 2:**
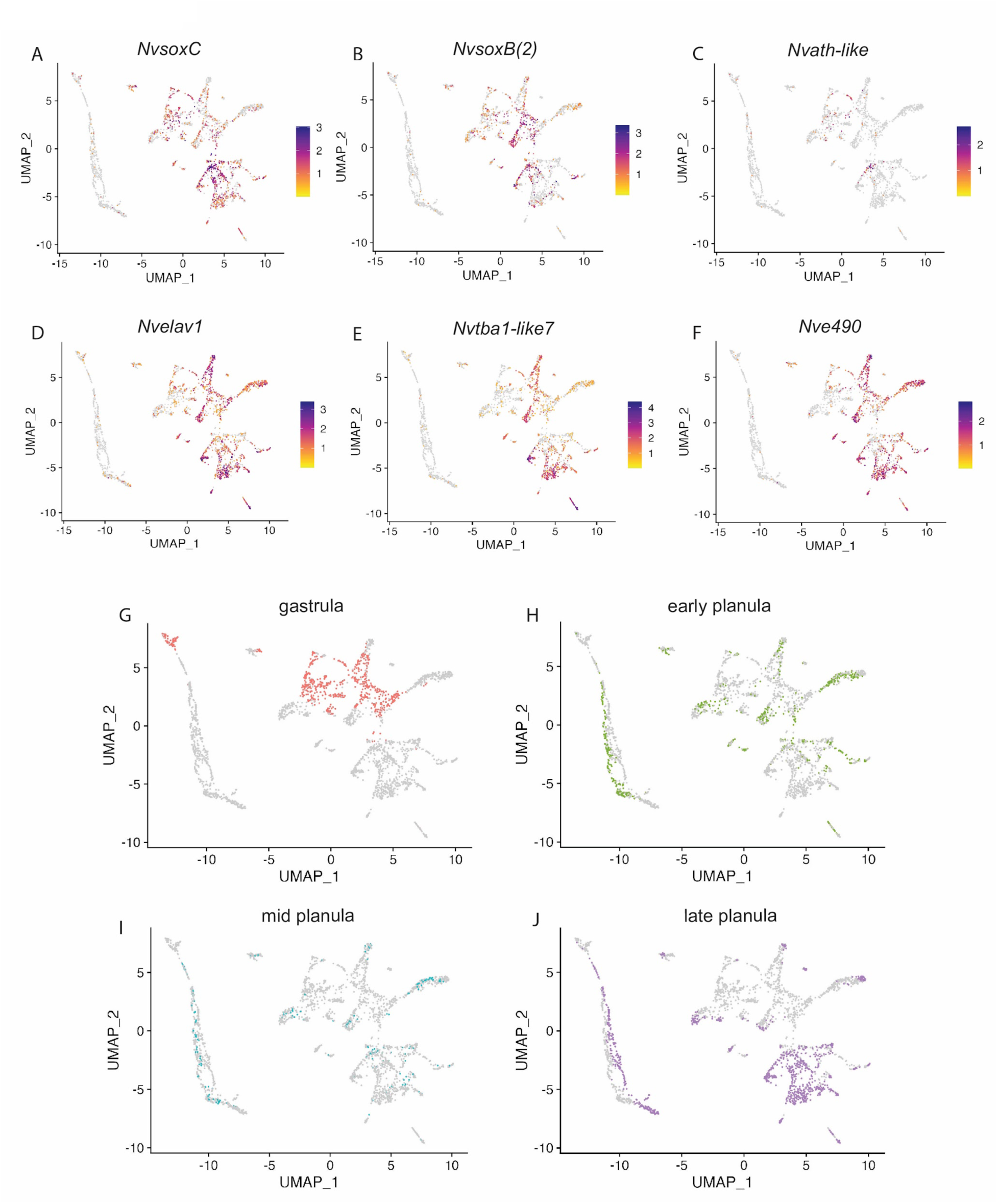
Expression of neural markers in the *NvashA* subset. (A-F) UMAPs highlighting the spatial expression of known early and late neural markers in the *NvashA* subset. (G-J) Individual UMAPs of the *NvashA* subset from Figure 4D, split to highlight the time-point from which each cell was obtained. (G) *NvashA* positive cells obtained from gastrula stage embryos are highlighted in red and are found in early neural (neuron.gast-pl) and cnidocyte (cnidocyte.gast and cnidocyte) clusters, as well as gland cells. Cells from early planula (H, green), mid planula (I, blue), and late planula (J, purple) stage animals are found in all but one of the cell types identified (except cnidocyte.gast).

**Supplemental Figure 3:**
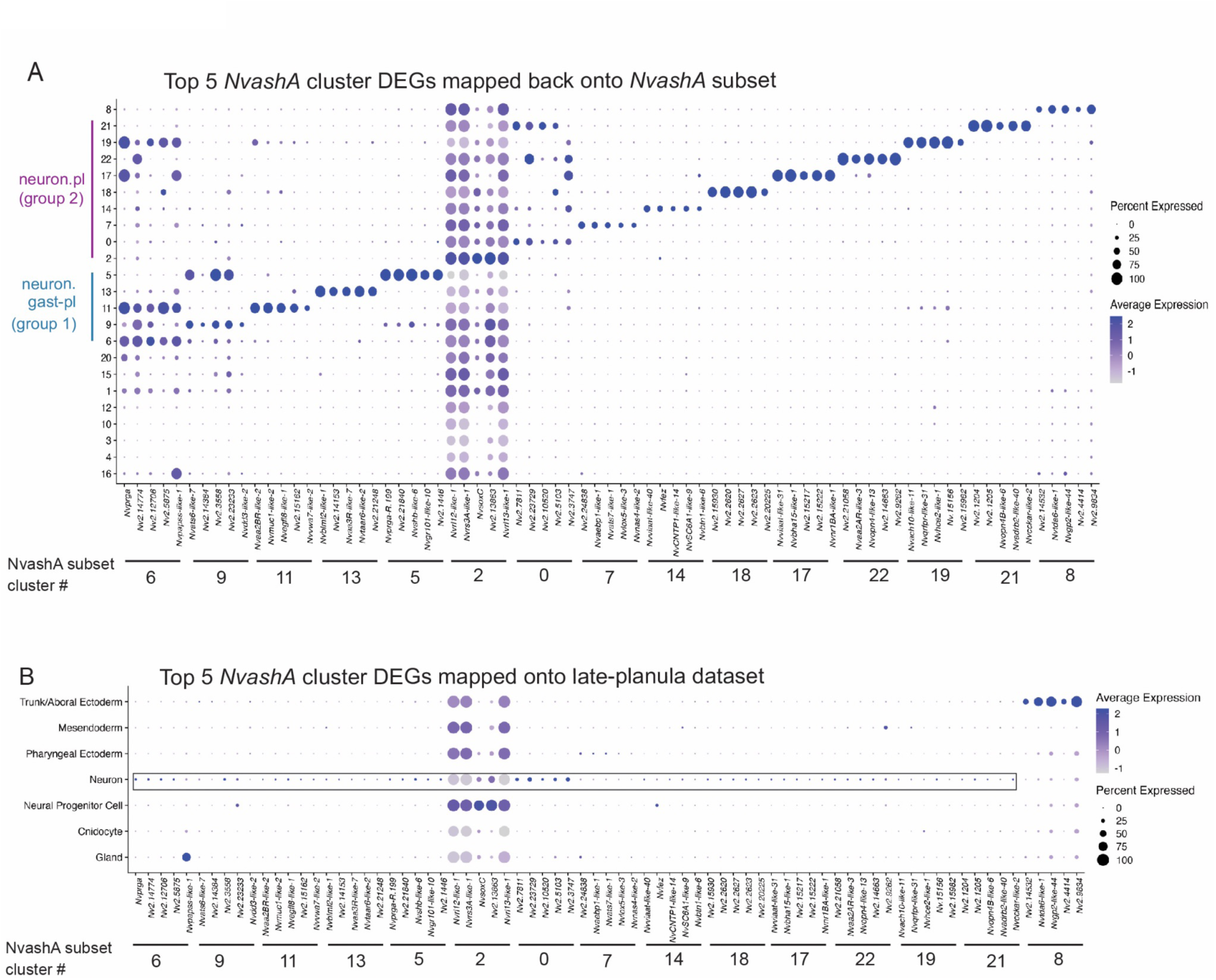
Use of differential gene expression analysis to identify novel potential cluster markers. (A) Dot plot representation of the top 5 DEGs identified for each neural cluster, mapped back onto the *NvashA* subset, to identify genes that are restricted to their respective cluster (and thus could represent good cluster marker genes). (B) Dot plot representation of the same genes in (A), mapped onto the late-planula dataset to demonstrate the neural specificity of the DEGs from the neural *NvashA* subset clusters. The majority of neural cluster markers were restricted to neuronal and neuroglandular clusters except markers from cluster 8.

**Supplemental Figure 4:**
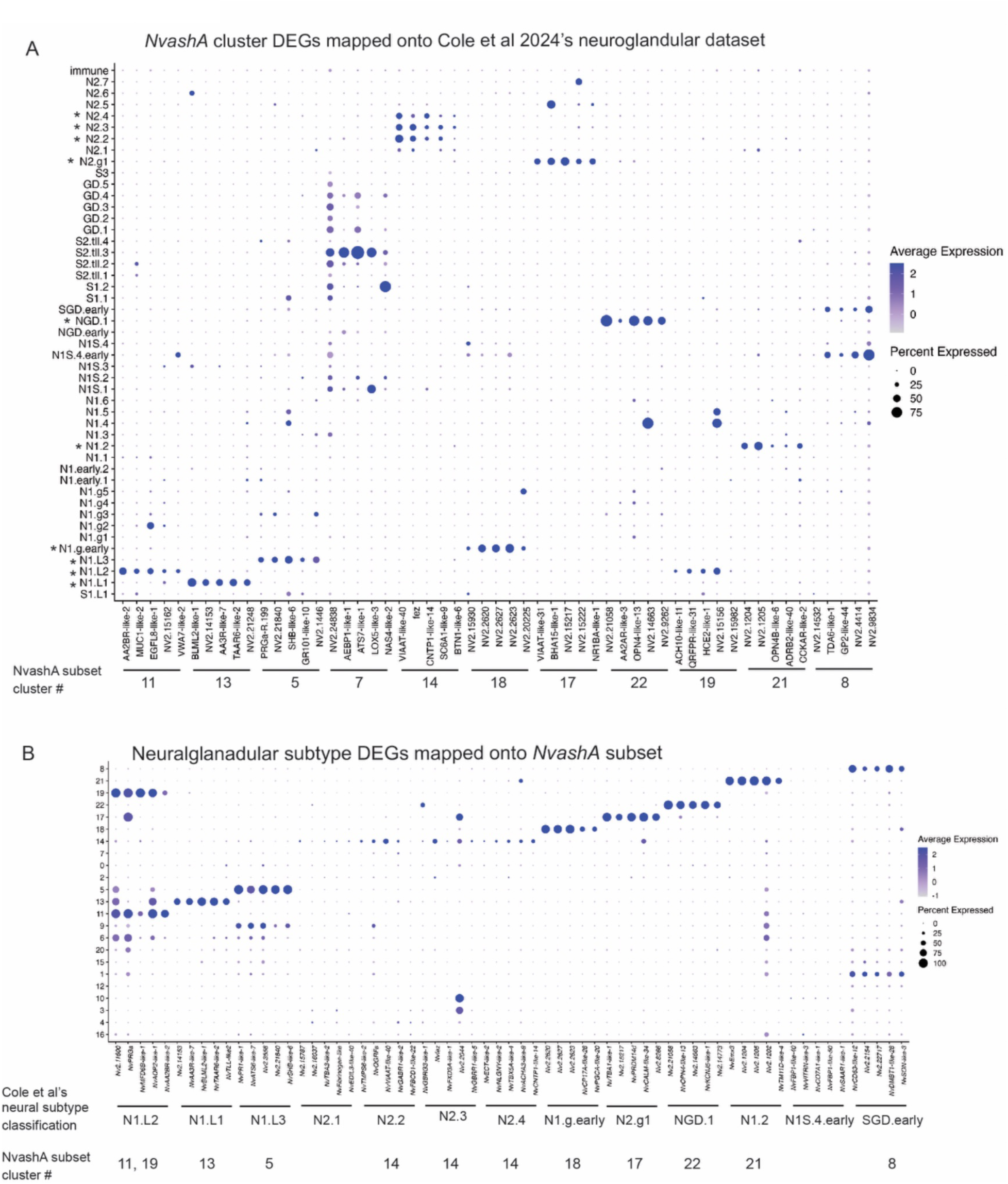
A subset of known neuroglandular subtypes are represented in *NvashA*-expressing cells through the late planula stage. (A) Dotplot illustrating the expression of the top DEGs from neural clusters within the *NvashA* subset across the neuroglandular dataset published by Cole et al. (2024). Clusters with restricted expression of their top DEGs to a known neuroglandular subtype were renamed to reflect that subtype (see Figure 1C). (B) Dotplot representation of the expression of the top 5 DEGs from the neuroglandular clusters/subtypes identified to be present in our *NvashA* subset in (A), reciprocally mapped onto our *NvashA* subset to confirm potential subtype identity of our clusters.

**Supplemental Figure 5:**
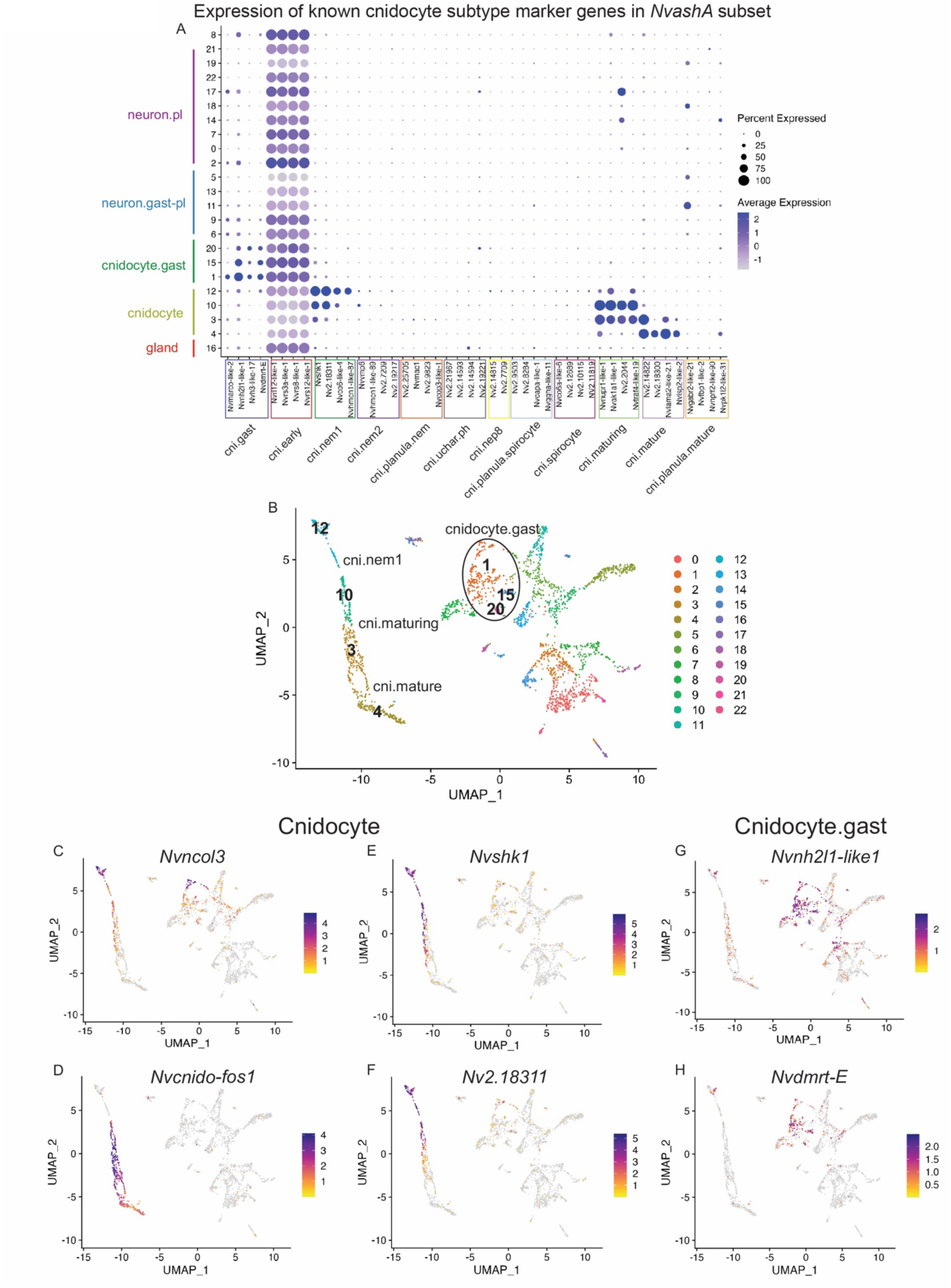
A subset of cnidocyte subtype genes are detected in *NvashA* expressing cells through the late planula stage. (A) Dotplot representation of the expression of published cnidocyte subtype markers (Steger et al, 2022) across *NvashA* subset clusters. (B) UMAP labeled with the clusters annotated as cnidocyte.gast, cni.nem1, cni.maturing, and cni.mature within the *NvashA* subset. (C-H) UMAPs showing the spatial expression of genes utilized to annotate (C-F) cnidocyte and (G,H) cnidocyte.gast clusters within the NvashA subset.

**Supplemental Figure 6:**
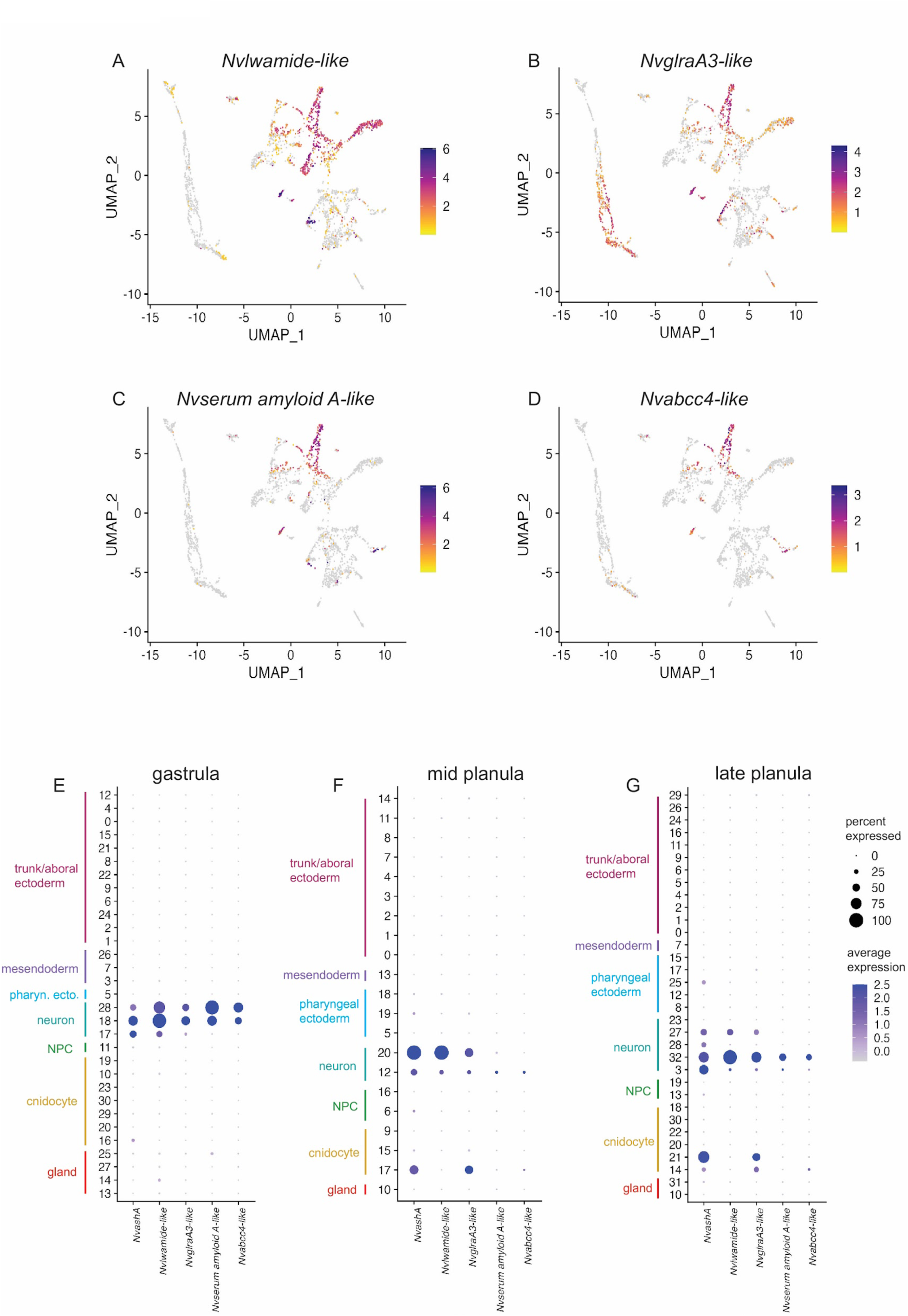
Known *NvashA* target genes are expressed in early- and late-derived cell clusters within the *NvashA* subset. (A-D) UMAP visualizations of *NvashA* subset showing the expression of the known *NvashA* targets (A) *NvLWamide-like*, (B) *NvglraA3-like*, (C) *Nvserum amyloid A-like*, and (D) *Nvabcc4-like*. (E-G) Dotplot representations of the expression of known *NvashA* target genes at (E) gastrula, (F) mid-planula, and (G) late-planula stages, demonstrating their continued expression throughout the developmental stages examined in this study.

**Supplemental Figure 7:**
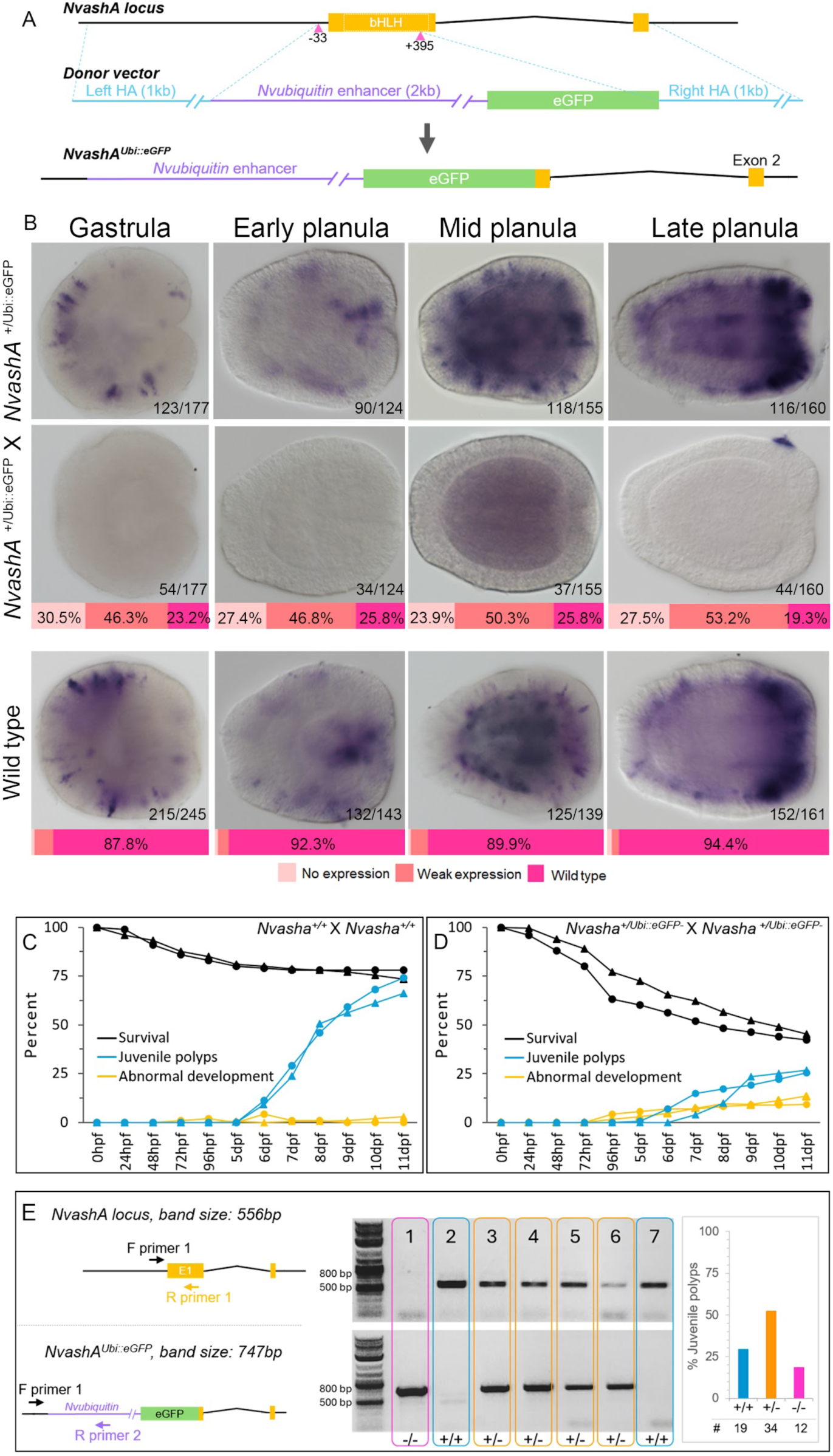
Generation and characterization of *NvashA* mutants. (A) Schematic of the CRISPR/Cas9 knock-in strategy of *Nvubiquitin::eGFP* into the *Nvasha* locus. Magenta triangles indicate PAM site locations. (B) *NvashA* expression in offspring from heterozygous *NvashA^+/Ubi::eGFp-^* mutant crosses (top), and offspring from wild type *NvashA^+/+^* crosses (bottom). In all images the oral pole is positioned to the right. (C-D) Percent of embryos that survive, develop into juvenile polyps, or have visually abnormal development over the first 11 days of development in offspring derived from (C) *NvashA^+/+^* X *NvashA^+/+^* and (D) *NvashA*^Ubi::eGFP+/-^ X *NvashA*^Ubi::eGFP+/-^ crosses (n=2 per cross, shown by circular and triangular markers). (E) Genotyping strategy for *NvashA*^Ubi::eGFP+/-^X *NvashA*^Ubi::eGFP+/-^ offspring using two primer sets (left), and representative image of genotyping PCR results for seven juvenile polyps, 11dpf (middle). At 11dpf, the genotypes of juvenile polyps from *NvashA*^Ubi::eGFP+/-^X *NvashA*^Ubi::eGFP+/-^ crosses follow Mendelian ratios for the *NvashA* mutant allele.

